# Wireless implantable sensor for non-invasive, longitudinal quantification of axial strain across rodent long bone defects

**DOI:** 10.1101/142778

**Authors:** Brett S. Klosterhoff, Keat Ghee Ong, Laxminarayanan Krishnan, Kevin M. Hetzendorfer, Young-Hui Chang, Mark G. Allen, Robert E. Guldberg, Nick J. Willett

## Abstract

Bone development, maintenance, and regeneration are remarkably sensitive to mechanical cues. Consequently, mechanical stimulation has long been sought as a putative target to promote endogenous healing after fracture. Given the transient nature of bone repair, tissue-level mechanical cues evolve rapidly over time after injury and are challenging to measure non-invasively. The objective of this work was to develop and characterize an implantable strain sensor for non-invasive monitoring of axial strain across a rodent femoral defect during functional activity. Herein, we present the design, characterization, and *in vivo* demonstration of the device’s capabilities for quantitatively interrogating physiological dynamic strains during bone regeneration. *Ex vivo* experimental characterization of the device showed that it exceeded the technical requirements for sensitivity, signal resolution, and electromechanical stability. The digital telemetry minimized power consumption, enabling long-term intermittent data collection. Devices were implanted in a rat 6 mm femoral segmental defect model and after three days, data were acquired wirelessly during ambulation and synchronized to corresponding radiographic videos, validating the ability of the sensor to non-invasively measure strain in real-time. Lastly, *in vivo* strain measurements were utilized in a finite element model to estimate the strain distribution within the defect region. Together, these data indicate the sensor is a promising technology to quantify local tissue mechanics in a specimen specific manner, facilitating more detailed investigations into the role of the mechanical environment in dynamic skeletal healing and remodeling.

## Introduction

Fracture healing is a dynamic physiological process requiring rapid and highly-coordinated morphogenesis of numerous cell populations to restore functional bone tissue. While the majority of the 12 million annual fractures in the United States heal without complications, 5-10% of fractures do not heal in a timely fashion and require lengthy clinical interventions involving multiple surgeries before function is restored [1–4]. To address this unmet clinical need, much research has been devoted to understanding mechanisms of fracture non-union and to developing novel therapeutic strategies to stimulate bone repair. As the primary load bearing tissue in the human body, bone development, maintenance, and regeneration are remarkably sensitive to mechanical cues [5–7]. Numerous studies have identified the critical role of local mechanical cues in tissue differentiation, formation, and functional restoration of bone defects [8–16]. Due to these findings, mechanical stimulation has long been sought after as a putative target to stimulate endogenous bone healing mechanisms.

Before moving to costly and resource intensive studies in large animals, the rodent femoral defect model has emerged as a primary pre-clinical test bed to evaluate novel therapeutics—including drugs, biologics, and scaffolds—for the treatment of load-bearing bone defects [17,18]. In this model, various fixation systems have been utilized with varying levels of load-sharing across the defect region. Ultimately, the mechanical environment experienced locally by cells in the defect milieu regulates the healing response [19]. Given the transient nature of bone repair, these tissue-level mechanical cues evolve rapidly with time after injury and are highly dependent on the specific injury model and treatment under investigation. However, the biomechanical environment in these models has rarely been analyzed and the multi-scale role of mechanical stimuli in the observed healing response remains poorly understood.

To elucidate the role of mechanical stimuli in skeletal healing, there is a need to quantify the mechanical environment experienced by the healing tissue during routine *in vivo* activities. However, standard techniques for quantitatively assessing the mechanical environment in pre-clinical animal models are limited to external fixation systems periodically affixed to mechanical loading instruments which impart a prescribed load stimulus to the defect [14,20,21], or computational image-based finite element (FE) simulations based on estimated mechanical boundary conditions [12]. The boundary conditions applied to such models are typically based on broad assumptions because they are challenging to measure non-invasively and change throughout the progression of healing. Consequently, neither technique directly captures loading patterns due to functional activities such as walking. Recognizing these limitations, we reasoned that the ability to directly and longitudinally measure the mechanical environment during fracture healing would enable quantification of the mechanical cues produced in specific bone defect models, and provide a more detailed understanding of mechanical stimuli that can promote or impair functional healing of skeletal injuries.

Sustained technological advancements in microelectronics and short-range wireless communication systems have rapidly refined biomedical sensor technologies, thus motivating research to explore implantable sensor based approaches to monitor an array of clinical diseases [22–25]. In particular, the introduction of small and inexpensive passive and active telemetry systems show promise for rapid deployment into pre-clinical models. The potential of implantable sensors to longitudinally monitor important environmental cues during regenerative processes in pre-clinical models could aid in the rational design and evaluation of novel therapeutics and advance understanding of the fundamental principles of mechanobiology [26]. To achieve this, we set out to engineer an implantable strain sensor— with a sufficiently small footprint to be utilized in the rodent femoral defect model—that can wirelessly transmit real time measurements of mechanical strain across a bone defect to a computer. To obtain accurate measurements an implantable sensing device should meet a number of critical functional requirements including:

1. Sufficient sensitivity and signal resolution to detect relevant changes in the parameter of interest (e.g. mechanical strain)
2. Limits of detection well outside the physiological dynamic range
3. Stable electromechanical characteristics throughout implantation under repeated functional loading and submersion in bodily fluids
4. Sufficient signal strength for wireless transmission through bodily tissues
5. Programmability and adequate power source for flexible, long-term data acquisition
6. Biocompatibility
7. Compatibility with longitudinal imaging techniques (e.g. radiography)
8. Mechanical durability to withstand surgical deployment and animal activity
9. Adequate size envelope for implantation in pre-clinical rodent models
10. Minimal external data acquisition hardware for logging wireless measurements to a PC

The objective of this work was to develop and characterize an implantable strain sensor for non-invasive, real-time monitoring of axial strain across a rodent femoral defect due to functional activity. Extensive *ex vivo* and *in vivo* validations were conducted to evaluate the device to meet the aforementioned functional criteria. Herein, we present the design, characterization, and *in vivo* evaluation of the device’s capabilities for quantitatively interrogating functional biomechanics post-fracture. Additionally, *in vivo* data were utilized in a finite element model to estimate the tissue-level mechanical environment within the defect.

## Materials and Methods

### Strain sensor device design

The implantable device consisted primarily of two components: (1) an internal fixation plate instrumented with a strain sensor and (2) a stacked board processing unit enabling sensor functionality and wireless data transmission. The modular fixation plate used to stabilize the rat femoral defect, as reported in previous studies, consists of a polysulfone segment which acts as the fixator and two stainless steel plates which interface the polysulfone segment with each end of the femur (Figure 1) [27,28]. The radiolucent polysulfone segment was modified to accommodate a single-element 350 Ω micro strain gage (Vishay PG EA-06-125BZ-350/E, Raleigh, NC) adhered into a recessed pocket on the back side by a two-component epoxy (Vishay PG M-Bond AE-10) while permitting *in vivo* radiographic imaging of the healing defect. Insulated lead wires running to the processing unit were soldered to the strain gage pads, cannulated through medical grade silicone tubing, and routed into a recessed channel on the side of the plate. To protect the sensor from fluid infiltration and mechanical impingement during surgery, the wire channel and pocket containing the strain gage was potted with a biocompatible (ISO 10993) light-curing sealant (Dymax 1072-M, Torrington, CT) and shielded by thin (380μm thick) laser-cut polytetrafluoroethylene (PTFE) covers.

**Figure 1:**
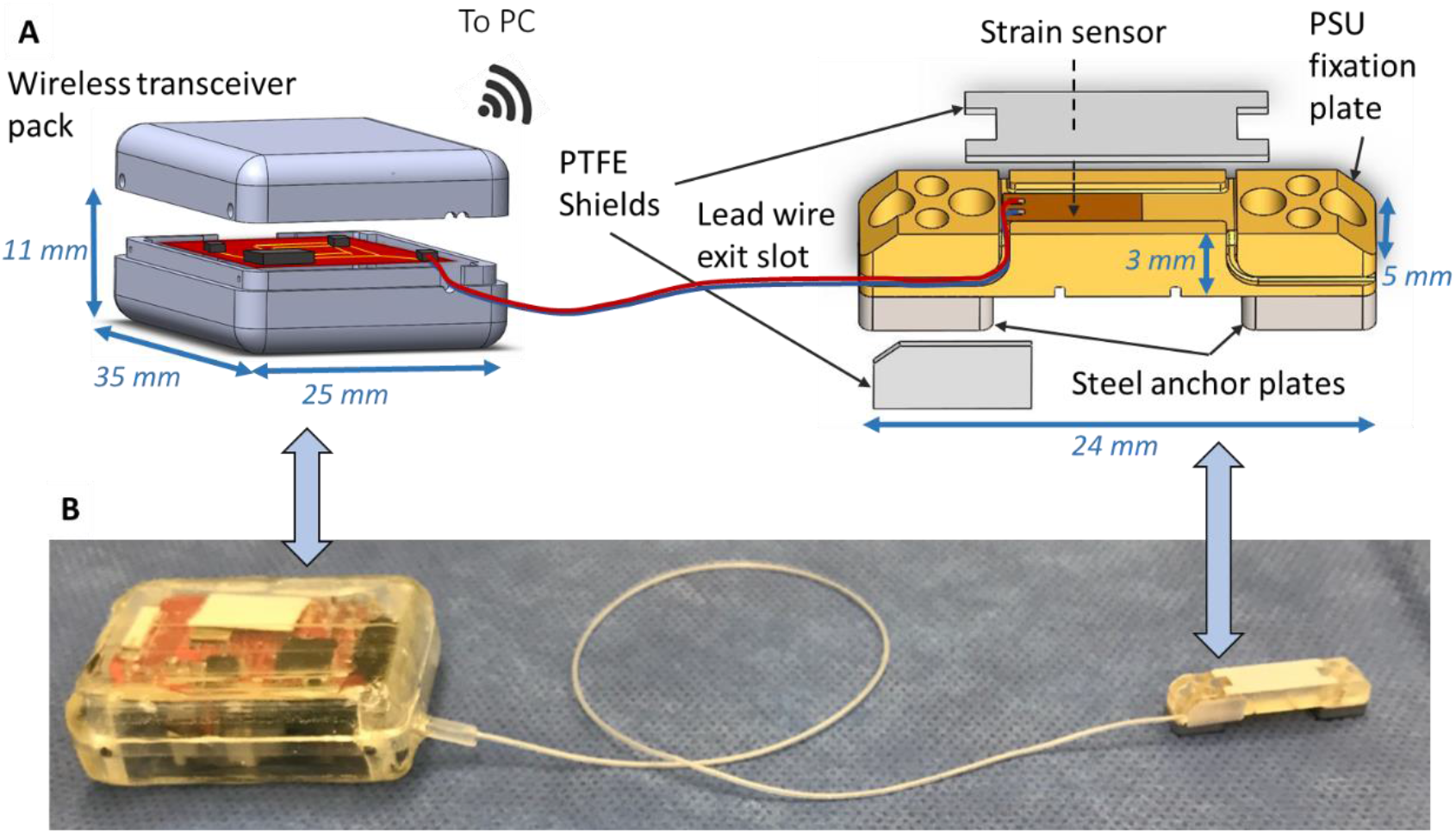
(a) Exploded view schematic of instrumented internal femoral fixation plate with sensor adhered in recessed pocket on plate and lead wires routed through machined slot to the transceiver pack mounted in the abdominal cavity. Note: plate and transceiver schematics are not to same scale (b) Photograph of instrumented plate before surgical implantation.

The stacked chip processing unit consisted of an active wireless network system (Texas Instruments EZ430-RF2500, Dallas, TX) featuring an ultra-low power microcontroller and 2.4 GHz RF transceiver. The network chip was interfaced with a custom signal conditioning circuit including a Wheatstone bridge, low pass filter, two-stage amplifier, voltage regulator, and 240 mAh lithium coin cell battery (Energizer CR-2032, St. Louis, MO) permitting approximately 33 total hours of active data transmission at 7-8 Hz. After connecting the circuit to the sensor lead wires, the entire stacked chip was housed in a custom 3D printed pack (Stratsys RGD720, Eden Prairie, MN) and encapsulated with the same biocompatible sealant used for the strain sensor. Wireless sensor data were acquired in real-time via a paired transceiver and USB plug-in mounted to a remote computer using a custom MATLAB graphical user interface (Mathworks, Natick, MA).

### Electromechanical characterization

The electromechanical characteristics of three instrumented fixation plates were evaluated *ex vivo* to assess the sensor’s sensitivity to detect strains due to physiological ambulatory loads and robustness to sustain long-term measurements in the *in vivo* environment. In order to simulate the eccentric bending loads placed on the internal fixation plates during functional loading, each end of the plates were attached to pre-drilled and tapped aluminum blocks approximating the length and cross-sectional area of the distal and proximal ends of the femur post-osteotomy using the same screws used to anchor the plate to the femur during surgery (Figure 2a). After assembly, a mechanical testing instrument (Electroforce 3220, TA Instruments, New Castle, DE) was used to apply cyclical compressive axial loads to the aluminum block-fixation plate construct. The constructs were pre-loaded to -2.5 N and tested for 25 cycles by a 0.5 Hz sinusoidal waveform to various magnitudes under displacement control, creating tensile local strains on the back surface of the fixation plate to which the strain sensor was adhered. Each test condition was repeated in triplicate for all three devices. The maximum displacement magnitude for each test was selected to achieve resultant axial load magnitudes of 16-20 N, which was the estimated peak axial force on the femur of a 250 g rat based on kinetic analysis of rodent gait [29]. In addition to testing the sensor with an empty gap between the two aluminum blocks, cylindrical defect surrogate materials of 40A polyurethane rubber and PTFE were placed to fill the gap between the blocks to mimic the stiffness of the bone defect region during different stages of healing, including the proliferating soft tissue callus stage and the eventual mineralization stage (Figure 2b; Rubber: elastic modulus = 4.766 ± 0.1153 MPa, axial stiffness = 19.26 ± 0.3646 N/mm; PTFE: elastic modulus = 319.3 ± 2.100 MPa, axial stiffness = 681.3±39.11 N/mm). Throughout testing, the local axial strain along the sensor region of the fixation plate was measured by laser extensometer (LX-500, resolution = 1μm, MTS, Eden Prairie, MN) while simultaneously recording the differential voltage output of the strain sensor to validate the measurements (Figure 2c).

**Figure 2:**
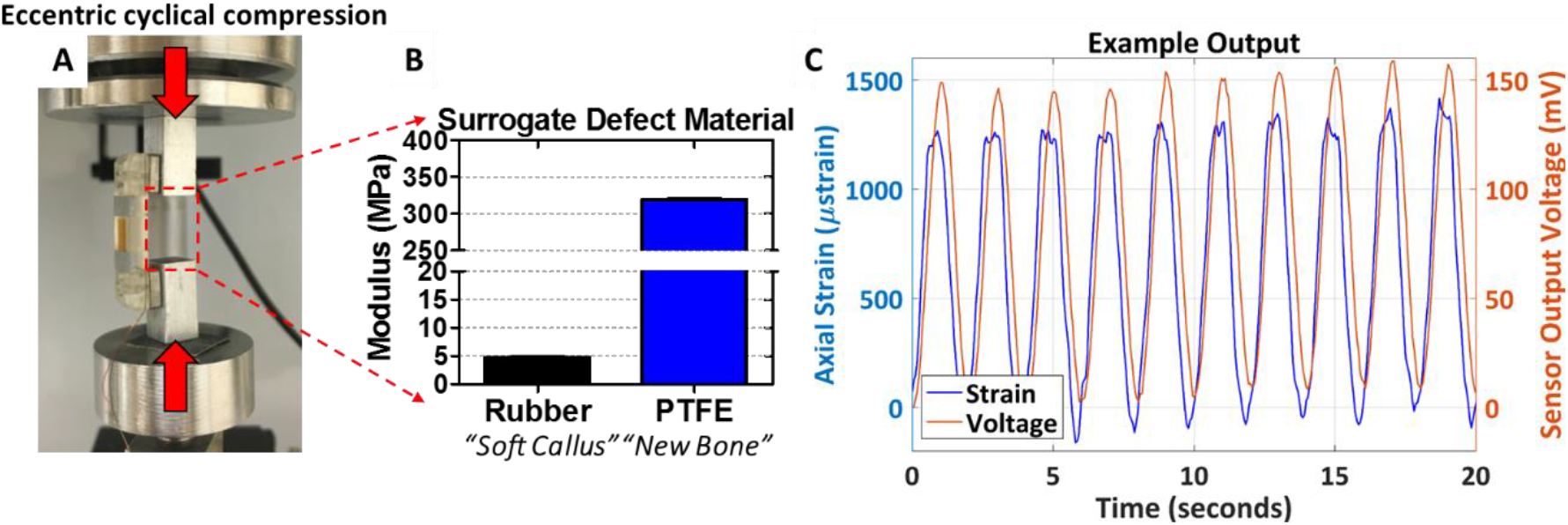
(a) Experimental set-up for eccentric cyclical compression testing of instrumented fixation plates. (b) Elastic moduli of surrogate defect materials, which are placed in the gap between the loading blocks to simulate the progression of mechanical properties during bone defect repair. (c) Example output during a cyclic test, where local strain in the sensor region as measured by laser extensometer is plotted alongside the corresponding voltage signal from the sensor.

### Fatigue testing

In order to assess the durability of the sensor to endure repeated flexural strains due to ambulatory loading, three plates were cyclically loaded in three-point bending to achieve the maximum anticipated local axial strain on the sensor *in vivo* (6800 με). Loads were applied for 10,000 cycles at 1 Hz while recording the output from the sensor.

### Biostability

To ensure the electromechanical characteristics of the sensor were stable under prolonged exposure to bodily fluids, instrumented fixation plates (n=2) were submerged in phosphate buffered saline (PBS) and placed in an oven at 37 °C for 4 weeks. Prior to submersion and at weekly intervals thereafter, the sensitivity of the strain sensor was quantified by loading the fixation plate in three-point bending over a range of physiological strain magnitudes on the sensor from 300 to 6800 με.

### Surgical procedures

All procedures were approved by the Georgia Institute of Technology Institutional Animal Care and Use Committee (IACUC protocol A14040). After arrival, rats were single housed for 1-2 weeks after procurement for acclimatization before experimental use, with unlimited access to food and water under a 12:12 hour light:dark cycle throughout the study. Unilateral 6 mm segmental defects were surgically created in the femurs of 8 month old male retired breeder Sprague Dawley rats weighing approximately 550-650 grams (n=2, CD, Charles River Labs, Wilmington, MA) under isofluorane anesthesia (1.5-2.5%) using a modification of previously established procedures to implant the transceiver pack in the abdominal cavity [30]. Briefly, the left femur was exposed from an anterior approach using blunt dissection. Next, the abdominal cavity was exposed by a 2 cm midline skin incision initiating 1 cm below the sternum (xiphoid) followed by an incision of the abdominal musculature (on the linea alba) and peritoneum, avoiding the sub-diaphragmatic organs. A keyhole incision was made through the abdominal wall superior to the left inguinal ligament and the fixation plate and associated wire were passed through the keyhole into the hindlimb compartment before positioning the transceiver pack into the abdominal cavity. The peritoneum and abdominal musculature were then sutured and the skin was closed with wound clips. The keyhole incision was carefully sutured with a loose loop around the traversing wire to prevent herniation while permitting translation of the wire during joint movement. The fixation plate was mounted to the anterolateral aspect of the femur using 4 screws and a critically sized 6 mm defect was created in the mid-diaphysis using an oscillating saw and left untreated. A subcutaneous injection of sustained-release buprenorphine was administered for analgesia prior to surgery.

### Wireless data acquisition during treadmill walking and high-speed radiography

One week prior to surgery, animals were trained to walk on a rat treadmill (Metabolic Modular Treadmill, Columbus Instruments, Columbus, OH) at slow ambulatory speeds ranging from 0.08-0.16 m/sec. A custom radiographic imaging system consisting of bi-plane X-ray generators and X-ray image intensifiers (Imaging Systems & Service, Inc., Painesville, OH) modified with the addition of high-speed digital video cameras (Xcitex XC-2M, Woburn, MA) was utilized to obtain high-speed images of the animal’s skeleton during walking periods. Three days after surgery, animals were imaged (100 frames/s; Camera 1 – 42 kV, 80 mA, 5 ms exposure; Camera 2 – 40 kV, 80 mA, 5 ms exposure) while walking on the treadmill at 0.1 m/sec. Throughout all treadmill activities, strain sensor readings were recorded wirelessly at 7-8 Hz.

### Finite element analysis

To ascertain the mechanical environment within the defect region during ambulation on the treadmill, a simplified FE model was constructed (ABAQUS CAE 6.14, Dassault Systèmes, Waltham, MA) and *in vivo* strain sensor data were utilized as boundary conditions. Briefly, a simplistic representation of the femur was generated by tying both ends of a cylinder representing the defect zone tissue (diameter = 5 mm; height = 6mm) to rectangular blocks representing the distal and proximal ends of the intact femur cortical bone (length = width = 6 mm; height = 17 mm). The fixation plate assembly was tied to the femur and an axial load was applied along the long-axis of the femur. Linear, elastic, homogeneous, isotropic mechanical properties were assigned to each component of the model as shown in Table 4, with defect tissue properties approximating early stage granulation tissue (Elastic modulus = 0.99 MPa, Poisson’s ratio = 0.33) [31]. The magnitude of the femoral load was varied so that the axial strain values of elements on the fixation plate surface where the sensor was adhered matched the 95^th^ percentile strain amplitude recorded during the *in vivo* treadmill acquisition period (approximately 5500 με). The 95^th^ percentile strain amplitude range was selected because it was the largest recorded strain magnitude range corresponding to 30-50 gait cycles in both animals, which is a sufficient number of loading stimuli to induce a physiological response [32–34]. The resultant axial load through the femur and maximum principal strain within the defect tissue were analyzed as the primary outcomes of the model.

### Statistical analysis

All data are presented as mean ± standard deviation unless otherwise stated. Multiple group comparisons were assessed by analysis of variance (ANOVA), with pairwise comparisons analyzed using Tukey’s post-hoc test (Graphpad Prism 7, San Diego, CA). A one-sample two-tailed Student’s t-test was utilized to evaluate fatigue testing results. A p-value < 0.05 was defined as a statistically measurable difference.

## 3. Results

### Electromechanical characterization

The results of off-axis electromechanical characterization are summarized in Figure 3 and Table 1. Sensors exhibited high linearity (Table 1; r^2^=0.993 ± 0.017, p<0.0001 all tests) and distinct sensor outputs for each test condition, (Figure 3a-b; Empty = 29.65 ± 1.51 mV/N, Rubber = 21.69 ± 1.45 mV/N, PTFE = 3.17 ± 1.24 mV/N, p<0.001 all comparisons), demonstrating the sensor had sufficient sensitivity to detect changes in axial strain on the fixation plate under loading at different healing stages (with different stiffness surrogate defect materials). Taking into account op-amp gain and excitation voltage used for empty and rubber surrogate mechanical testing, the overall sensitivity of the device was 0.287±0.035 μV/V/με. Sensitivity under PTFE surrogate test conditions could not be computed as local plate strains were at or below the resolution of the laser extensometer and could not be reliably measured, indicating the sensor was more sensitive than the extensometer.

**Figure 3:**
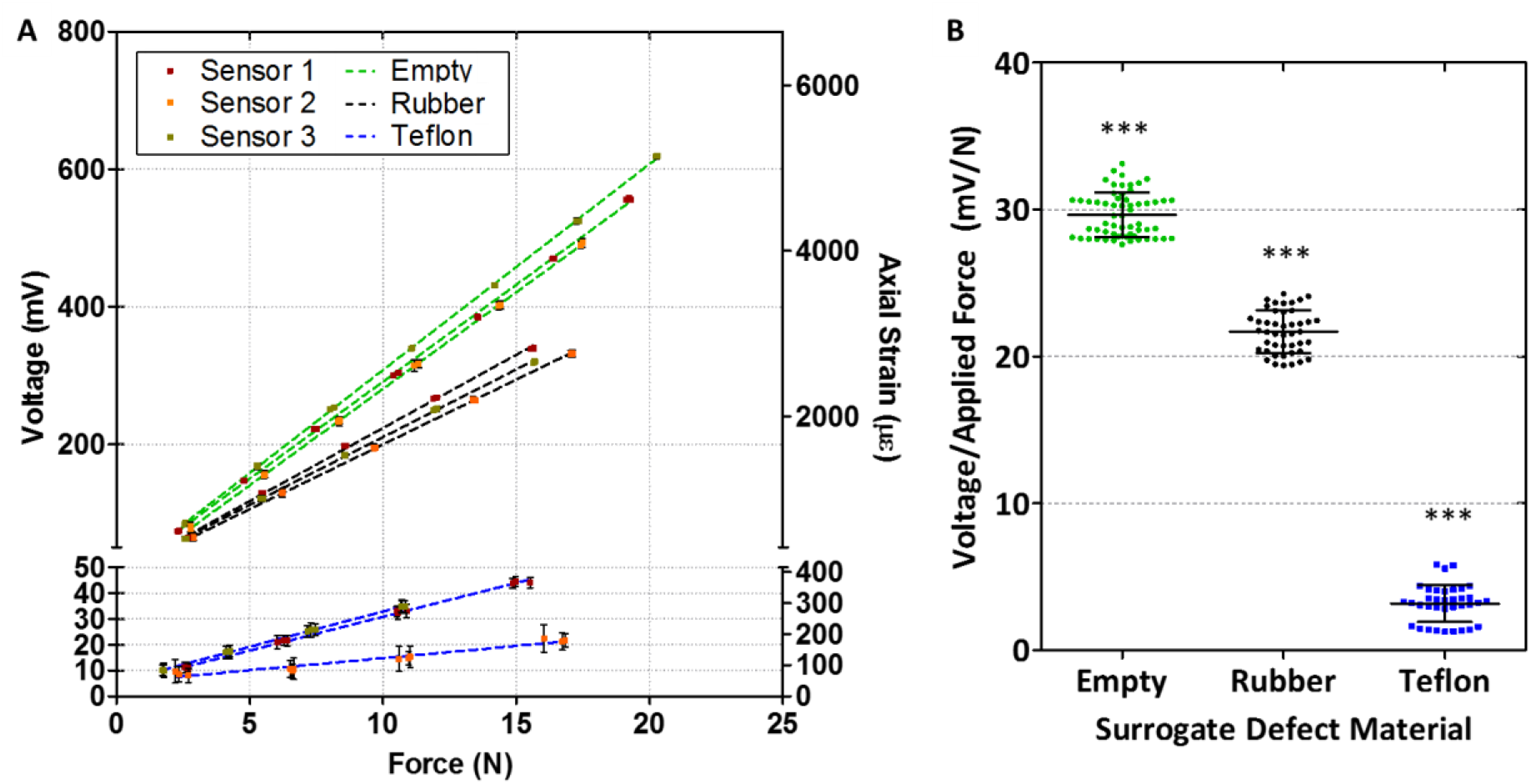
(a) Sensor output plots for a range of physiological load magnitudes. The color of the dot represents a different sensor and the color of the line represents a different defect surrogate material. Cyclical tests were repeated in triplicate for each loading case, and error bars depicting standard deviation are included on all data points. (b) Normalizing the voltage output of the sensor by applied force demonstrates the sensor is able to discern changes in the stiffness of the defect region, and therefore appears promising to detect progression of bone repair under physiological load conditions, *** p<0.001, ANOVA, all comparisons.

**Table 1:**
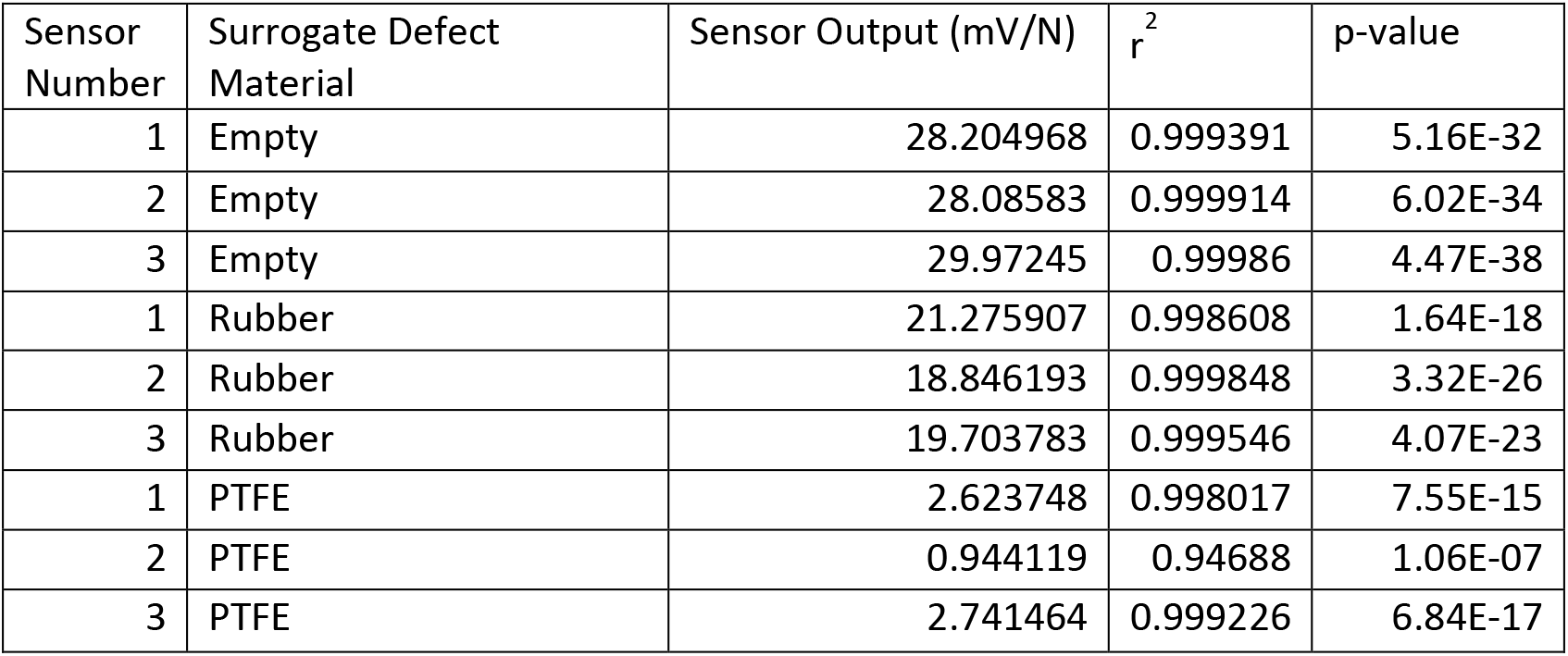
Sensor output and linear regression results for electromechanical characterization testing in Figure 2.

To quantify the lower limit of detection of the device, we computed the signal-to-noise ratio (SNR) of the mechanical testing output and performed linear regression as a function of axial strain on the fixation plate (Figure 4, Table 2). A limit of detection cut-off criteria of 20 dB (corresponding to a signal amplitude-to-noise ratio of 10-1) was applied and linear regression computed that 300 με was the minimum detectable strain amplitude on the fixation plate that would be utilized in analysis of *in vivo* testing. Strains of this magnitude were only observed in PTFE surrogate defect testing, suggesting the sensor possessed sufficient resolution to accurately characterize the strain due to functional loading until and potentially after complete and robust bridging of the entire bone defect occurred.

**Figure 4:**
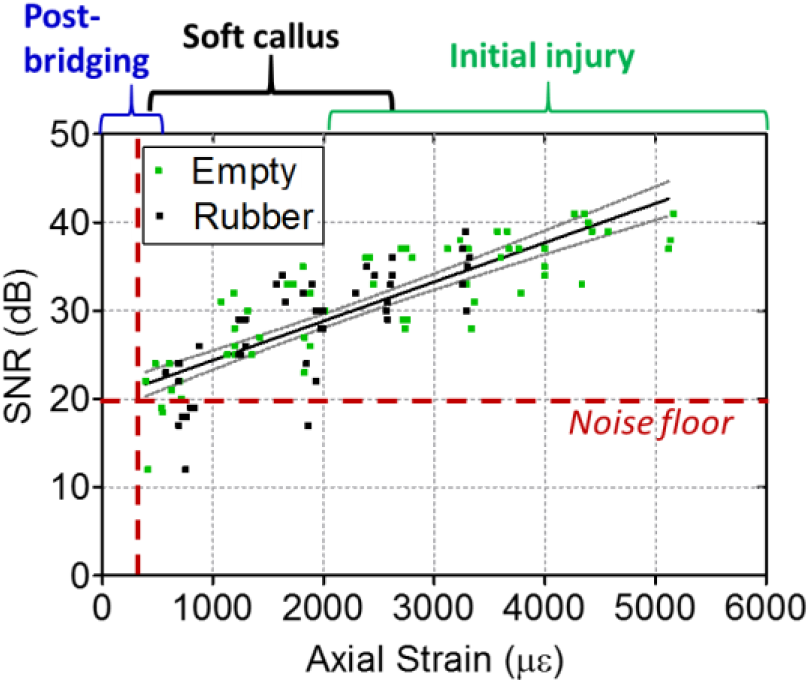
Regression of signal to noise ratio (SNR) of sensor output versus the local axial strain of the sensor region as measured by laser extensometer. Employing a limit of detection cut-off criteria of 20 dB (corresponding to a signal amplitude-to-noise ratio of 10-1) indicates the sensor can reliably detect plate strains as low as 300 με, suggesting the sensor possessed sufficient resolution to obtain measurements until and potentially after robust bridging of the bone defect occurred.

**Table 2:** Linear regression of 95% confidence interval (CI) for SNR vs. strain depicted in Figure 3. Conservatively employing the lower bounds of the CI to compute the SNR gives 19.52 dB for 300 με, which consequently is defined as the limit of detection.

| Linear Regression 95% Confidence Intervals |  |
| --- | --- |
| Slope (dB/$\mu\epsilon$) | 0.003818 to 0.005067 |
| Y-intercept (dB) | 18.37 to 21.55 |
| $r^2$ | 0.6618 |

**Table 3:**
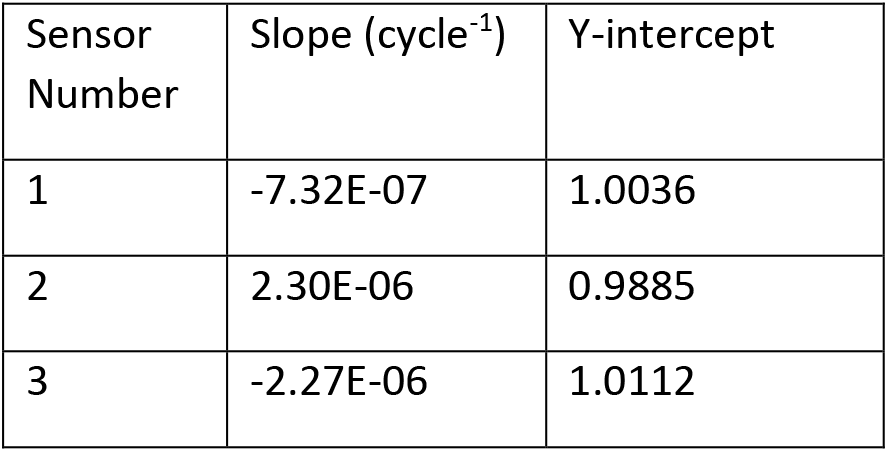
Linear regression results for sensor fatigue testing.

| Sensor Number | Slope (cycle<sup>-1</sup>) | Y-intercept |
| --- | --- | --- |
| 1 | -7.32E-07 | 1.0036 |
| 2 | 2.30E-06 | 0.9885 |
| 3 | -2.27E-06 | 1.0112 |

**Table 4:** Mechanical properties of components utilized in the definition of the FE model. Biological tissue mechanical properties are obtained from [31,38]

| Component | Material | Elastic modulus (MPa) | Poisson's ratio |
| --- | --- | --- | --- |
| Femur | Cortical bone | 18000 ref. [38] | 0.3 |
| Defect tissue | Granulation tissue | 0.99 ref. [31] | 0.33 |
| Fixation plate | Polysulfone | 2500 | 0.37 |
| Fixator screw plates | Stainless steel | 200000 | 0.33 |

### Fatigue testing

The normalized output of sensors (n=3) throughout 10,000 cycles was analyzed to determine if repeated deformation at the maximum anticipated strain *in vivo* would alter its sensitivity (Figure 5a). Throughout testing, the response of the sensors was stable within ± 4%. The slope of the regression lines (Table 2; -2.32×10^−7^ ± 2.32 ×10^−6^) was not significantly different from 0 (p=0.877), demonstrating the sensor was sufficiently durable to maintain a constant sensitivity under high cycle peak physiological loads.

**Figure 5:**
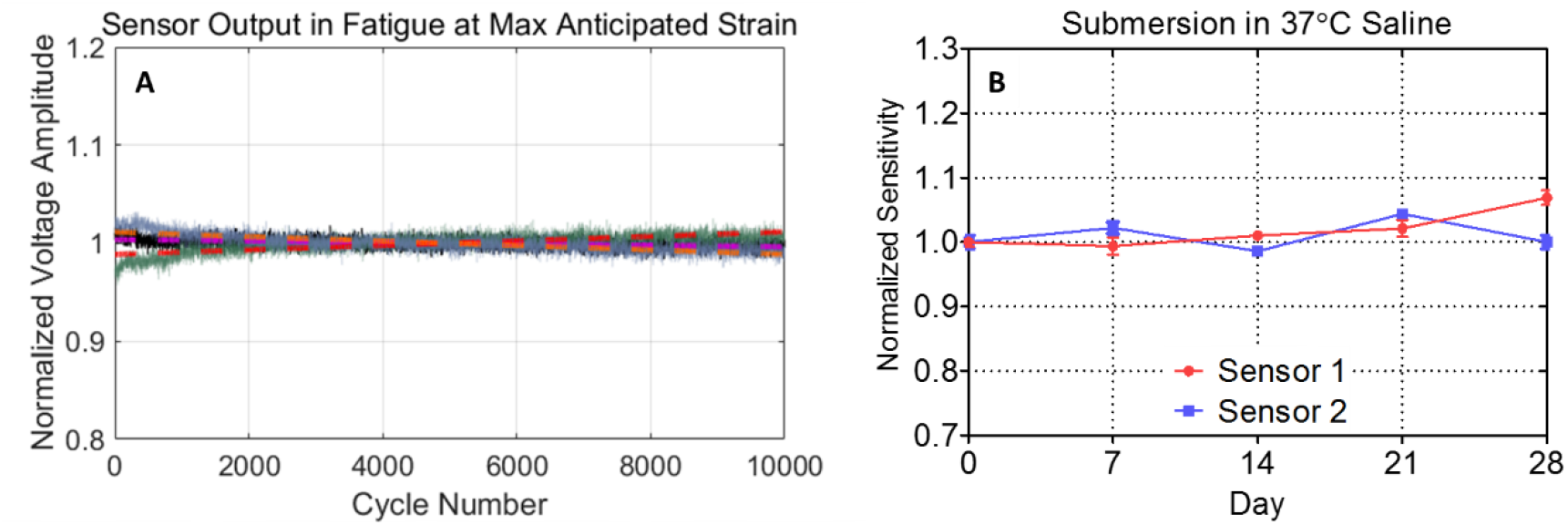
(a) Fatigue testing of the devices (n=3) for 10,000 cycles at maximum anticipated physiological strain. The normalized raw outputs for each device is shown, and their respective regression lines are depicted by the bright dotted lines. Throughout 10,000 cycles the outputs were stable within +/-4 percent and the resultant slope was not significantly different than zero (p=0.877). (b) Instrumented fixation plates were submerged in saline maintained at body temperature for 4 weeks and sensitivity was evaluated by mechanical testing at weekly intervals. The output was stable within 7% throughout the test and sensitivity plots for all time-points remained highly linear (r^2^ range = 0.9938-0.9988).

### Biostability

Throughout submersion in PBS at body temperature, the sensitivity of the sensor was stable within 7% of the value measured prior to submersion (Figure 5b). High linearity was maintained throughout testing trials (r^2^ range = 0.9938-0.9988). These data indicate that the sensor can sustainably provide precise measurements under long term physiological conditions.

### Wireless strain data acquisition and radiographic imaging of rodent gait after surgical implantation and creation of femoral defect

After implantation of two devices in untreated 6 mm femoral defects, strain data were recorded wirelessly by a nearby laptop. Stable wireless acquisition was maintained up to a distance of approximately 20 feet, with further increasing distance resulting in lost data packets and thereby reducing sampling frequency. The presence of metal tables or objects in the transmitter line-of-sight was determined to reduce the transmission range to as low as 5 feet, depending on the spatial configuration. Three days after surgery, the rats were walked on a treadmill at a slow speed of 0.1 m/sec. The belt speed was selected to produce a gait cycle of about 1 Hz, which was sufficiently slow to mitigate the risk of aliasing the gait cycle while recording strain data at 7-8 Hz. The logged sensor data were synchronized to 6 second videos of the animal recorded by high-speed radiographic videos from two different angles (Supplementary Videos 1 and 2). The synchronized videos demonstrated the sensor signal coincides with the phase and frequency of the operated limb gait cycle, with lower strains being measured when the leg is lifted and rapid increases in strain observed when the leg is planted.

Absolute strains on each plate were computed from the *in vivo* voltage signal based on the calibrated sensitivity determined by electromechanical characterization of each device prior to implantation. To account for changes in the zero strain set-point which occur while surgically anchoring the fixation plate to the femur, the voltage corresponding to zero static strain was estimated by the median voltage signal while the animal sat in its cage prior to the treadmill walking period, which was nearly constant due to the minimal motion of the animal. During the videos, peak strains of up to 3290 με were observed during loading phases, whereas strain reversal relaxed the static flexural strain on the plate up to -3140 με while the leg was lifted (Figure 6).

**Figure 6:**
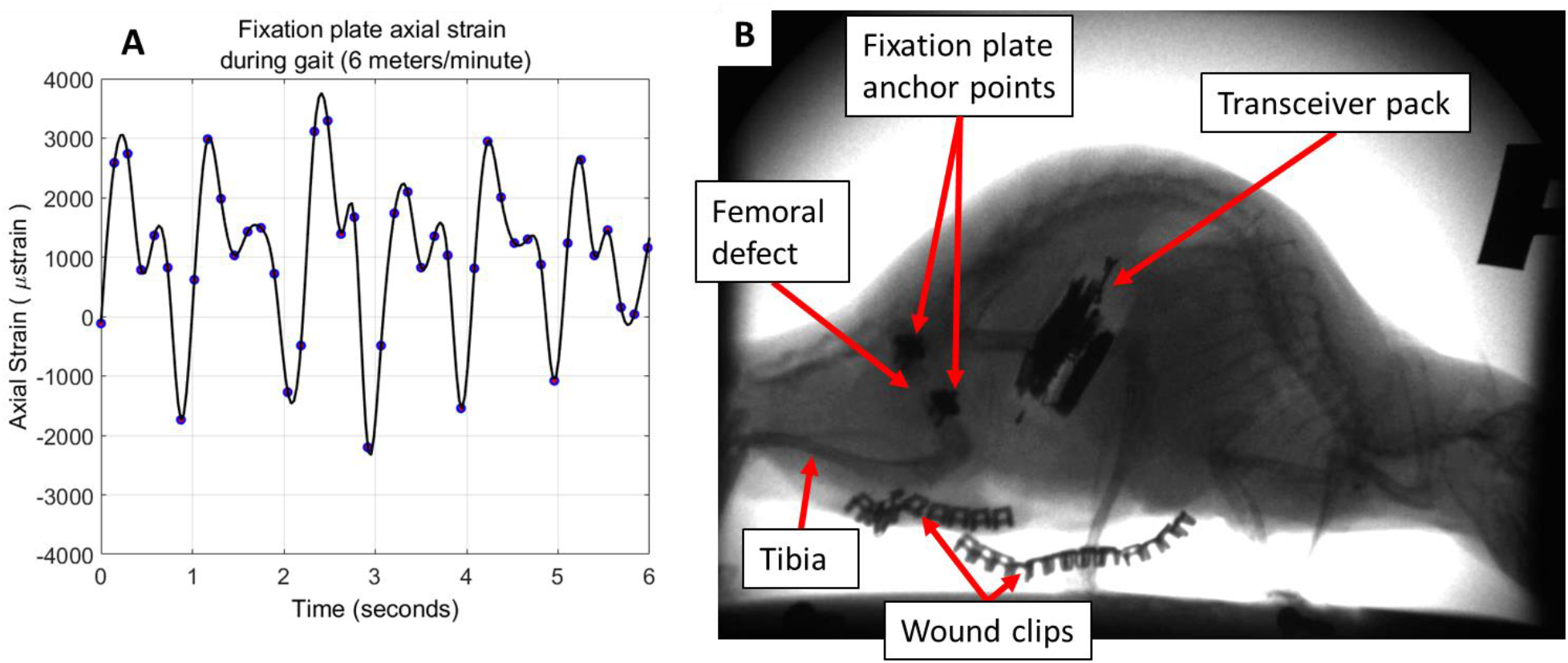
(a) Representative *in vivo* strain versus time measurement recorded wirelessly during ambulation on a treadmill three days after the creation of a 6 mm segmental femoral defect. Actual data points are depicted as circles with a spline curve-fit illustrated by the black line. (b) During data acquisition, high-speed radiographic videos were acquired by two x-ray cameras mounted at different angles. The videos were synchronized with the recorded sensor output to validate the ability of the sensor to non-invasively quantify functional strains in real-time (see Supplementary Videos 1 and 2). Radiopaque objects including the stainless steel components which anchor the femoral fixation plate, the abdominally implanted transceiver circuit pack, and the incision wound clips are labelled.

### Fixation plate strain analysis

The distribution of dynamic strain cycles experienced by the fixation plate during the entire treadmill walking period for both animals were quantified using a peak analysis to compute strain amplitudes between adjacent local minima and maxima in the recorded sensor signals (MATLAB). The load history revealed a number of similarities between the two animals. Histograms of the strain amplitudes indicate a skewed distribution with approximately 35% of the total strain cycles falling in relatively low strains between 300 to 1000 με (Figure 7). The incidence of strain cycles between 1000 to 5000 με were observed to be nearly constant for both implants. The median and 95^th^ percentile strain amplitude were computed to be 1929 με and 5543 με, and 1889 με and 6041 με for each implant respectively.

**Figure 7:**
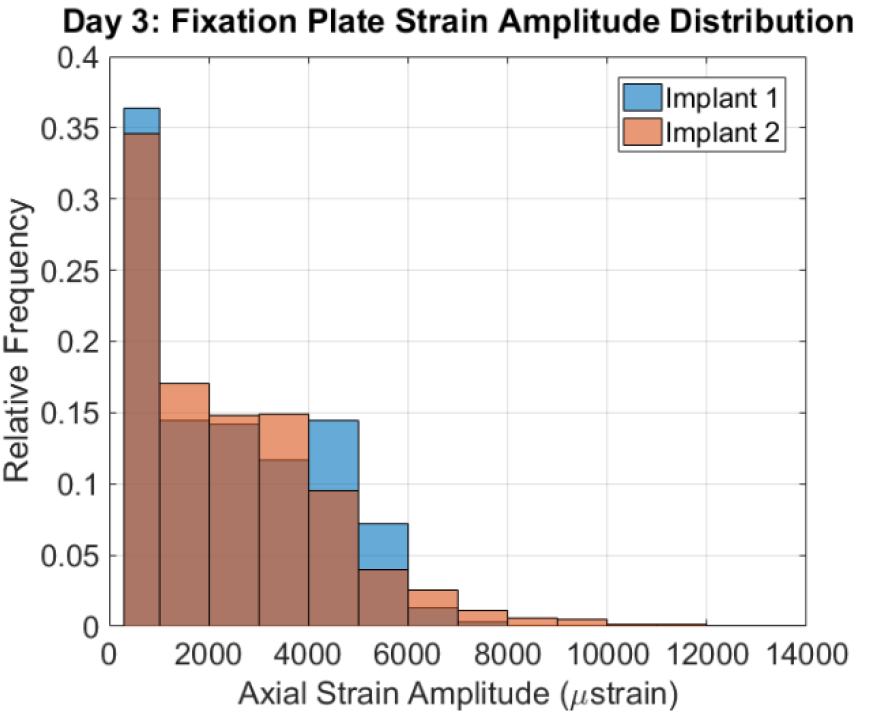
Strain amplitude distributions during ambulation. The median and 95^th^ percentile strain amplitude were computed to be 1929 με and 5543 με, and 1889 με and 6041 με for each implant respectively.

### Defect tissue finite element analysis

A simplified FE model of the femur and fixation plate was constructed to estimate the mechanical environment within the defect zone. The resultant axial load on the femur was varied to match the local axial strain recorded on the fixation plate region where the strain sensor was adhered. The resultant axial load was computed to be 26.3 N, corresponding to 3.6 times the animals’ bodyweight (Figure 8a). Compressive deformations were observed throughout the defect region, with a gradient of increasing strain as the distance from the neutral axis of the fixation plate increased. The minimum principal strain of the elements comprising the defect tissue was 8.85 ± 1.95%, with the majority of the defect tissue undergoing compressive strains between 6 and 12% (Figure 8b).

**Figure 8:**
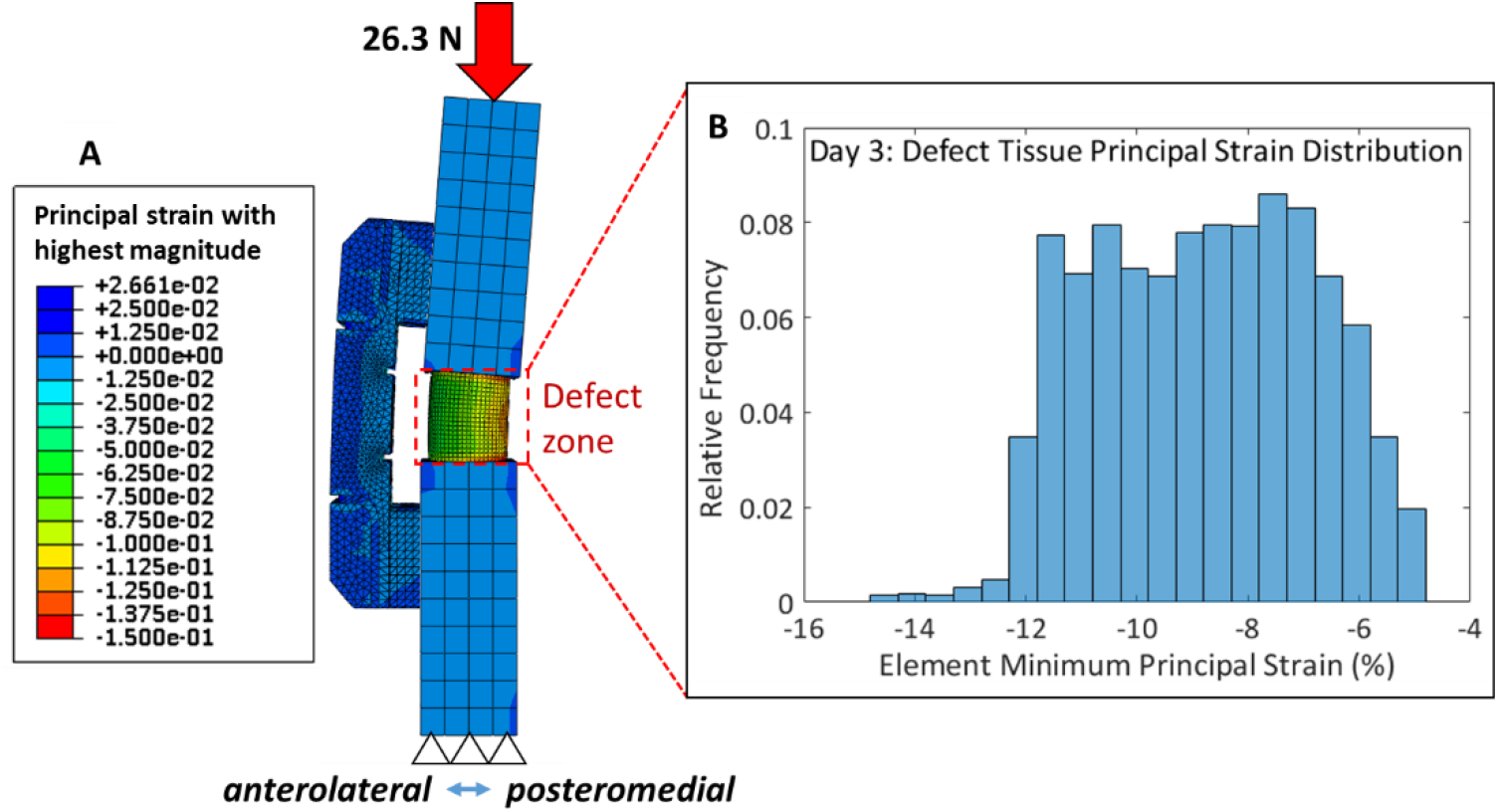
(a) Simplified FE model of fixation plate mounted to femur, with analysis focused on a cylindrical “defect zone” 6 mm in length. The axial load on the femur (26.3 N) was back-calculated to match the 95^th^ percentile local axial strain in the sensor region measured during ambulation. Linear, elastic homogenous mechanical properties were assigned for all components and are tabulated in Table 4. (b) The average minimum principal strain within the defect zone was found to be 8.85 ± 1.95%.

## 4. Discussion

The mechanical environment within healing tissue evolves rapidly with time and is challenging to quantify longitudinally in a non-invasive fashion. The aim of this study was to develop, characterize and evaluate the ability of an internal fixation plate with an integrated wireless strain sensor to non-invasively quantify dynamic axial strains across a rodent bone defect during functional activity. Key technical criteria were defined and evaluated through *ex vivo* and *in vivo* experiments. In summary, the data indicate the sensor possesses sufficient sensitivity to detect progressive changes in physiological strain as healing occurs. Additionally, the lower limit of detection was sufficient to obtain strain measurements until robust bridging of the entire defect occurs. The sensor was proven to withstand 10,000 cycles at the maximum *in vivo* strain with no change in sensitivity, indicating the dynamic range of the sensor encompasses the relevant physiological range. Submersion testing demonstrated the sensor is packaged and sealed in a manner which maintains electromechanical stability for sustained physiological measurements. *In vivo* implantation demonstrated the device could be surgically implanted into a rat femoral segmental defect model and successfully transmit data wirelessly to a nearby computer. Animals tolerated the abdominal implant well throughout the study. Strain measurements were acquired in real-time while animals walked on a treadmill and the data were validated by synchronizing to high-speed radiographic videos recorded simultaneously. High-speed x-ray imaging was a desirable validation technique for the sensor because it allowed for direct and accurate visualization of the position of the long bones during the gait cycle, overcoming movement artifacts caused by the large amount of soft tissue mass surrounding the hindlimb which limits the accuracy of standard optical imaging techniques [35].

The primary contribution of the reported device is the capability to longitudinally acquire quantitative measurements describing the *in vivo* evolution of the mechanical environment during bone healing. Perren et al. were the first to develop a predictive theory of mechanical boundary conditions that determine the differentiation of skeletal tissues after fracture [8]. Since then, many studies have utilized a variety of experimental and computational techniques to elucidate tissue-level mechanical conditions that enhance or impair bone repair [9–16]. While these studies have refined our understanding of the mechanobiological regulation of fracture healing, the majority of the studies rely on non-physiological mechanical stimuli [14,15,20,21] or assumed boundary conditions for computational analysis that are not determined from the specific model under investigation [12]. A key limitation in the field is the technical challenge of longitudinally and non-invasively measuring mechanical conditions in *in vivo* models of skeletal healing [36,37]. By incorporating a wireless sensor, the mechanical boundary conditions of the defect can be quantified during physiological activities without disrupting the normal activity of the animal to obtain the measurement. The non-destructive measurement approach maximizes the amount of data acquired from a single animal, thereby reducing the number of animals needed to fully characterize the healing process and enabling specimen-specific analyses over multiple time points. Here we report a wireless device that has the capability to facilitate flexible, long-term data acquisition at discrete time points. The fully digital telemetry approach with an integrated ultra-low-power (50 μA) sleep mode permits the 33 hours of total power budget for active data transmission to be allocated over extended periods of time by programming the microcontroller for intermittent transmission. In our initial validation study, we selected a measurement protocol to obtain the majority of *in vivo* data within the first 72 hours post-surgery. In future studies, the measurement periods can easily be reprogrammed to extend the device over weeks and months.

The device also has some limitations. First, the single-element strain sensor is only capable of quantifying mechanical strain in one direction; in the configuration reported here it was along the axis of the femur. Wehner and colleagues performed kinetic analysis of rodent gait to demonstrate this direction is the primary load trajectory during ambulation [29]. In their analysis, bending moments in the mid-diaphysis were also observed in all anatomical planes. Based on this analysis and our own observations in the eccentric loading tests and FE model, the sensor was positioned in the optimal region to detect the primary mode of deformation. In the segmental defect model, the addition of the fixation plate naturally produces an eccentric axial load on the plate and out-of-plane bending, resulting in peak local tensile strains on the sensor while the defect zone undergoes compression. The plate cross-section is designed so the bending moment of inertia is threefold higher to in-plane bending that cannot be detected by the sensor, so its contribution to the overall mechanical environment is less significant. Second, the single-element design is sensitive to temperature differences between the hindlimb and abdomen. However, the *in vivo* conditions are essentially isothermal between the two regions as confirmed experimentally by the negligible drift in the baseline voltage of the sensor throughout implantation. Future improvements to address the aforementioned limitations could include a full-bridge strain sensor that is sufficiently small for the internal fixation plate, but commercial sensors satisfying these size constraints do not currently exist. We are actively pursuing custom full-bridge strain sensors fabricated via microelectromechanical systems (MEMS) manufacturing processes for this application.

*In vivo* strain measurements acquired three-days after surgery during walking on a treadmill demonstrate the fixation plate and defect tissue are subjected to a range of strain magnitudes, with primarily lower magnitudes but a significant number of higher magnitude loading events. A highly-simplified finite element model of the femoral defect and fixation plate was constructed to obtain an initial estimate of the mechanical environment within the tissue. Back-calculating the axial load on the femur to match the experimental strain sensor data provided validated boundary conditions for the model. The resultant axial load, 26.3 N, corresponded to 3.6 times the bodyweight of the animals. Wehner et al computed the peak axial force through the femur to reach up to 7 times bodyweight in naive rats [29]. The relatively small twofold discrepancy is likely due to gait deficits in the animals after undergoing a significant segmental defect surgery resulting in substantial damage to the hindlimb musculature as well as buprenorphine pain management three days prior to data acquisition. Noticeable favoring of the operated leg was apparent at this time point. Another contributing factor could be the walking speed. In our model, a relatively slow walking speed of 0.1 m/sec was chosen, whereas the gait speed in Wehner and colleague’s analysis was unreported.

The minimum principal compressive strain in the defect tissue was determined to be primarily between 6 and 12%. Mechanoregulation theories developed in non-critically sized fracture healing models have indicated mechanical stimuli in this range promote endochondral ossification [10,11,14]. The model defect investigated here is critically sized and functional healing does not occur without the addition of an osteogenic stimulus such as recombinant human bone morphogenetic protein 2 (rhBMP-2). Instead, a highly heterogeneous non-union tissue is formed consisting primarily of fibrous tissue after 12 weeks [30]. Thus, direct comparisons between the untreated critically sized defect with healing fracture mechanoregulation theories is not necessarily pertinent, but a mix of cartilage and bone has been observed in defects treated with 0.5 or 2.5 μg rhBMP-2 delivered in an alginate-based hydrogel system in this model three weeks after surgery [13]. Nonetheless, the initial data provide a valuable estimate of actual deformations in the defect due to physiological activity shortly after injury. In the future, computational analyses can be refined by incorporating *in vivo* micro-CT-based tissue geometries and mechanical properties of both the femur and defect tissue and will enhance our ability to interrogate the local mechanical environment within the regenerating tissue over time.

To conclude, we report the characterization and initial *in vivo* data obtained by a novel implantable wireless strain sensor in a pre-clinical model of bone repair. The device is capable of measuring axial mechanical strain during physiological activities and met key technical criteria outlined in the introduction. The technological underpinnings are broadly applicable to the mechanical characterization of therapeutics in diaphyseal fracture or defect models. This sensor platform provides a novel means to longitudinally characterize local tissue mechanics in a specimen specific manner, enabling more detailed investigations into the role of the mechanical environment in skeletal healing.

## Acknowledgements

This work was supported by a research partnership between Children’s Healthcare of Atlanta and the Georgia Institute of Technology and grants from the National Institutes of Health (NIH R21 AR066322; NIH R01 AR069297) and the National Science Foundation (NSF CMMI-1400065). BSK was supported by the Cell and Tissue Engineering NIH Biotechnology Training Grant (T32-GM008433) and the National Science Foundation Graduate Research Fellowship Program (DGE-1650044).

## References

[1] Pollak, A., and Watkins-Castillo, S., 2014, Fracture Trends - The Burden of Musculoskeletal Diseases in the United States.

[2] Zura, R., Xiong, Z., Einhorn, T., Watson, J. T., Ostrum, R. F., Prayson, M. J., Della Rocca, G. J., Mehta, S., McKinley, T., Wang, Z., and Steen, R. G., 2016, “Epidemiology of Fracture Nonunion in 18 Human Bones,” JAMA Surg., 151(11), p. e162775.

[3] Tarchala, M., Harvey, E. J., and Barralet, J., 2016, “Biomaterial-Stabilized Soft Tissue Healing for Healing of Critical-Sized Bone Defects: the Masquelet Technique.,” Adv. Healthc. Mater.

[4] Ilizarov, G. A., 1989, “The tension-stress effect on the genesis and growth of tissues. Part I. The influence of stability of fixation and soft-tissue preservation.,” Clin. Orthop. Relat. Res., (238), pp. 249–81.

[5] Frost, H. M., 2001, “From Wolff’s law to the Utah paradigm: Insights about bone physiology and its clinical applications,” Anat. Rec., 262(4), pp. 398–419.

[6] Rot, C., Stern, T., Blecher, R., Friesem, B., and Zelzer, E., 2014, “A Mechanical Jack-like Mechanism Drives Spontaneous Fracture Healing in Neonatal Mice,” Dev. Cell, 31(2), pp. 159–170.

[7] Gerstenfeld, L. C., Cullinane, D. M., Barnes, G. L., Graves, D. T., and Einhorn, T. A., 2003, “Fracture healing as a post-natal developmental process: Molecular, spatial, and temporal aspects of its regulation,” J. Cell. Biochem., 88(5), pp. 873–884.

[8] Perren, S. M., 1979, “Physical and biological aspects of fracture healing with special reference to internal fixation.,” Clin. Orthop. Relat. Res., (138), pp. 175–96.

[9] Goodship, A. E., and Kenwright, J., 1985, “The influence of induced micromovement upon the healing of experimental tibial fractures.,” J. Bone Joint Surg. Br., 67(4), pp. 650–5.

[10] Claes, L. E., Claes, L. E., Heigele, C. a, Heigele, C. a, Neidlinger-Wilke, C., Neidlinger-Wilke, C., Kaspar, D., Kaspar, D., Seidl, W., Seidl, W., Margevicius, K. J., Margevicius, K. J., Augat, P., and Augat, P., 1998, “Effects of mechanical factors on the fracture healing process.,” Clin. Orthop. Relat. Res., 355S, pp. S132–47.

[11] Claes, L.., and Heigele, C.., 1999, “Magnitudes of local stress and strain along bony surfaces predict the course and type of fracture healing,” J. Biomech., 32(3), pp. 255–266.

[12] Boerckel, J. D., Kolambkar, Y. M., Stevens, H. Y., Lin, A. S. P., Dupont, K. M., and Guldberg, R. E., 2012, “Effects of in vivo mechanical loading on large bone defect regeneration,” J. Orthop. Res., 30(7), pp. 1067–1075.

[13] Boerckel, J. D., Uhrig, B. A., Willett, N. J., Huebsch, N., and Guldberg, R. E., 2011, “Mechanical regulation of vascular growth and tissue regeneration in vivo,” Proc. Natl. Acad. Sci., 108(37), pp. E674–E680.

[14] Morgan, E. F., Salisbury Palomares, K. T., Gleason, R. E., Bellin, D. L., Chien, K. B., Unnikrishnan, G. U., and Leong, P. L., 2010, “Correlations between local strains and tissue phenotypes in an experimental model of skeletal healing.,” J. Biomech., 43(12), pp. 2418–24.

[15] Miller, G. J., Gerstenfeld, L. C., and Morgan, E. F., 2015, “Mechanical microenvironments and protein expression associated with formation of different skeletal tissues during bone healing.,” Biomech. Model. Mechanobiol.

[16] Betts, D. C., and Müller, R., 2014, “Mechanical regulation of bone regeneration: theories, models, and experiments.,” Front. Endocrinol. (Lausanne)., 5, p. 211.

[17] Histing, T., Garcia, P., Holstein, J. H., Klein, M., Matthys, R., Nuetzi, R., Steck, R., Laschke, M. W., Wehner, T., Bindl, R., Recknagel, S., Stuermer, E. K., Vollmar, B., Wildemann, B., Lienau, J., Willie, B., Peters, A., Ignatius, A., Pohlemann, T., Claes, L., and Menger, M. D., 2011, “Small animal bone healing models: standards, tips, and pitfalls results of a consensus meeting.,” Bone, 49(4), pp. 591–9.

[18] Horner, E. A., Kirkham, J., Wood, D., Curran, S., Smith, M., Thomson, B., and Yang, X. B., 2010, “Long bone defect models for tissue engineering applications: criteria for choice,” Tissue Eng. Part B. Rev., 16(2), pp. 263–271.

[19] Klosterhoff, B. S., Nagaraja, S., Dedania, J. J., Guldberg, R. E., and Willett, N. J., 2017, “Material and Mechanobiological Considerations for Bone Regeneration,” Materials and Devices for Bone Disorders, S. Bose, and A. Bandyopadhyay, eds., Academic Press, pp. 197–264.

[20] Wulsten, D., Glatt, V., Ellinghaus, A., Schmidt-Bleek, K., Petersen, A., Schell, H., Lienau, J., Sebald, W., Plöger, F., Seemann, P., and Duda, G. N., 2011, “Time kinetics of bone defect healing in response to BMP-2 and GDF-5 characterised by in vivo biomechanics.,” Eur. Cell. Mater., 21, pp.177–92.

[21] Schwarz, C., Wulsten, D., Ellinghaus, A., Lienau, J., Willie, B. M., and Duda, G. N., 2013, “Mechanical Load Modulates the Stimulatory Effect of BMP2 in a Rat Nonunion Model,” Tissue Eng. Part A, 19(1-2), pp. 247–254.

[22] Gibney, E., 2015, “The inside story on wearable electronics.,” Nature, 528(7580), pp. 26–8.

[23] McGilvray, K. C., Unal, E., Troyer, K. L., Santoni, B. G., Palmer, R. H., Easley, J. T., Demir, H. V., and Puttlitz, C. M., 2015, “Implantable microelectromechanical sensors for diagnostic monitoring and post-surgical prediction of bone fracture healing.,” J. Orthop. Res., 33(10), pp. 1439–46.

[24] Wise, K. D., 2007, “Integrated sensors, MEMS, and microsystems: Reflections on a fantastic voyage,” Sensors Actuators, A Phys., 136(1), pp. 39–50.

[25] Wachs, R. A., Ellstein, D., Drazan, J., Healey, C. P., Uhl, R. L., Connor, K. A., and Ledet, E. H., 2013, “Elementary Implantable Force Sensor: For Smart Orthopaedic Implants.,” Adv. Biosens. Bioelectron., 2(4).

[26] Klosterhoff, B. S., Tsang, M., She, D., Ong, K. G., Allen, M. G., Willett, N. J., and Guldberg, R. E., 2017, “Implantable Sensors for Regenerative Medicine,” J. Biomech. Eng., 139(2), p. 20806.

[27] Oest, M. E., Dupont, K. M., Kong, H.-J., Mooney, D. J., and Guldberg, R. E., 2007, “Quantitative assessment of scaffold and growth factor-mediated repair of critically sized bone defects,” J. Orthop. Res., 25(7), pp. 941–950.

[28] Kolambkar, Y. M., Dupont, K. M., Boerckel, J. D., Huebsch, N., Mooney, D. J., Hutmacher, D. W., and Guldberg, R. E., 2011, “An alginate-based hybrid system for growth factor delivery in the functional repair of large bone defects.,” Biomaterials, 32(1), pp. 65–74.

[29] Wehner, T., Wolfram, U., Henzler, T., Niemeyer, F., Claes, L., and Simon, U., 2010, “Internal forces and moments in the femur of the rat during gait.,” J. Biomech., 43(13), pp. 2473–9.

[30] Oest, M. E., Dupont, K. M., Kong, H.-J., Mooney, D. J., and Guldberg, R. E., 2007, “Quantitative assessment of scaffold and growth factor-mediated repair of critically sized bone defects.,” J. Orthop. Res., 25(7), pp. 941–50.

[31] Leong, P. L., and Morgan, E. F., 2008, “Measurement of fracture callus material properties via nanoindentation.,” Acta Biomater., 4(5), pp. 1569–75.

[32] Willie, B. M., Yang, X., Kelly, N. H., Han, J., Nair, T., Wright, T. M., van der Meulen, M. C. H., and Bostrom, M. P. G., 2010, “Cancellous Bone Osseointegration Is Enhanced by In Vivo Loading,” Tissue Eng. Part C Methods, 16(6), pp. 1399–1406.

[33] Ozcivici, E., Luu, Y. K., Adler, B., Qin, Y.-X., Rubin, J., Judex, S., and Rubin, C. T., 2010, “Mechanical signals as anabolic agents in bone,” Nat. Rev. Rheumatol., 6(1), pp. 50–59.

[34] Lanyon, L. E., 1984, “Functional strain as a determinant for bone remodeling,” Calcif. Tissue Int., 36(1), pp. S56–S61.

[35] Bauman, J. M., and Chang, Y.-H., 2010, “High-speed X-ray video demonstrates significant skin movement errors with standard optical kinematics during rat locomotion,” J. Neurosci. Methods, 186(1), pp. 18–24.

[36] Epari, D. R., Duda, G. N., and Thompson, M. S., 2010, “Mechanobiology of bone healing and regeneration: *in vivo* models,” Proc. Inst. Mech. Eng. Part H J. Eng. Med., 224(12), pp. 1543–1553.

[37] Claes, L. E., and Cunningham, J. L., 2009, “Monitoring the mechanical properties of healing bone.,” Clin. Orthop. Relat. Res., 467(8), pp. 1964–71.

[38] Ashman, R. B., Cowin, S. C., Van Buskirk, W. C., and Rice, J. C., 1984, “A continuous wave technique for the measurement of the elastic properties of cortical bone,” J. Biomech., 17(5), pp. 349–361.

